# Functional Connectivity of the Precuneus Reflects Effectiveness of Visual Restitution Training in Chronic Hemianopia

**DOI:** 10.1101/2020.05.14.050310

**Authors:** Hinke N. Halbertsma, Joris A. Elshout, Douwe P. Bergsma, David G. Norris, Frans W. Cornelissen, Albert V. van den Berg, Koen V. Haak

**Affiliations:** Laboratory for Experimental Ophthalmology, University Medical Center Groningen, Groningen, the Netherlands; Donders Institute for Brain, Cognition and Behaviour, Radboud University Medical Center, Nijmegen, the Netherlands

**Keywords:** Resting-State, Functional Connectivity, Visual Restitution Training, hemianopia, Precuneus, spatial attention

## Abstract

Visual field defects in chronic hemianopia can improve through visual restitution training, yet not all patients benefit equally from this long and exhaustive process. Here, we asked if resting-state functional connectivity prior to visual restitution could predict training success. In two training sessions of eight weeks each, 20 patients with chronic hemianopia performed a visual discrimination task by directing spatial attention towards stimuli presented in either hemifield, while suppressing eye movements. We examined two effects: a sensitivity change in the attended (trained) minus the unattended (control) hemifield (i.e., a training-specific improvement), and an overall improvement (i.e., a total change in sensitivity after both sessions). We then identified five visual resting-state networks and evaluated their functional connectivity in relation to both training effects. We found that the functional connectivity strength between the anterior Precuneus and the Occipital Pole Network was positively related to the attention modulated (i.e., training-specific) improvement. No such relationship was found for the overall improvement or for the other visual networks of interest. Our finding suggests that the anterior Precuneus plays a role in training-induced visual field improvements. The resting-state functional connectivity between the anterior Precuneus and the Occipital Pole Network may thus serve as an imaging-based biomarker that quantifies a patient’s potential capacity to direct spatial attention. This may help to identify hemianopia patients that are most likely to benefit from visual restitution training.

## 1. Introduction

Neurological rehabilitation for visual deficits (Raz and Levin, 2017) is increasingly recognised as an advantageous approach for improving the visual function of patients with chronic visual field defects (VFDs) such as homonymous hemianopia (i.e., one-sided cerebral blindness). Currently, three main rehabilitation approaches are used: substitutional, compensatory and restorative. The first two primarily aim for practical improvements of visual function of the sighted field with the help of visual aids or through improved effective visual scanning behaviour. Restorative approaches, in contrast, aim for a reduction of the VFD by increasing local sensitivity or reducing its perimetrical size. A substantial number of studies have reported functional improvements within the VFD measured using either ophthalmic, electrophysiological or psychophysical tests (Bergsma et al., 2012; Bergsma and Van Der Wildt, 2010; Bergsma et al., 2017; Elshout et al., 2018; Elshout et al., 2016; Huxlin and Cavanaugh, 2017; Julkunen et al., 2003; Kasten et al., 1998; Marshall et al., 2010; Mueller et al., 2007; Romano et al., 2008; Sahraie et al., 2006; Jobke et al., 2009) or neuroimaging (Julkunen et al., 2006; Marshall et al., 2010; Raemaekers et al., 2010).

The study presented here follows up on a controlled cross-over study in a cohort of chronic hemianopia patients that followed extensive visual restitution training (VRT) (Elshout et al., 2016). During this training, patients underwent a visual discrimination task that required directing spatial attention covertly to various locations in their visual field (VF). Following training, a significantly larger increase in VF sensitivity was found, at the group level, for the attended (i.e., trained) compared to the unattended (i.e., control) hemifield. Despite this positive training outcome at group level, the magnitude of this training effect was highly variable across patients. Such inter-subject variability in visual improvements following VRT has been reported before (Julkunen et al., 2003; Romano et al., 2008; Sabel, 2008), but has not been attributed to specific features of the training itself or the patient’s capacity to perform it. Because of the presence of an attentional component in the training paradigm of Elshout et al. (2016), their observation of inter-subject variability suggests that differences in individuals’ attentional capacities may underlie VRT success. This suggestion would be in line with the finding that the direction of attention towards a stimulus can improve visual restitution (Poggel et al., 2004), a capacity that may vary across individuals. The neural correlate(s) of such attentional capacity might be identified prior to training and thus could ultimately be developed into a biomarker for identifying patients that are most likely to profit from training. Such biomarker would be beneficial, as VRT protocols tend to be long and demand a lot from both patients and their care professionals.

Therefore, in the present study, we focussed on a possible *functional mechanism* that underlies VRT-induced improvement and explored this in the context of directing spatial attention. We did this by investigating the resting-state (RS) functional connectivity (FC) between brain regions of hemianopia patients, as examined by functional MRI. There are several reasons why exploring RS FC could be valuable in the context of neurological rehabilitation for visual deficits. First, RS FC is determined based on the temporal correlations between spontaneous signal fluctuations in spatially separated brain regions that occur when participants are not engaged in a specific task. For that reason, RS FC is particularly suitable for patients that cannot be visually stimulated in the same way as healthy controls, or for patients that have trouble with task-compliance. Secondly, the mere presence of a particular structural pathway does not necessarily imply that it is used functionally. Thirdly, RS FC enables identification of non-visual areas that are potentially relevant contributors to VRT-induced visual improvement, but do not possess a retinotopic organisation. Fourthly, RS FC can be assessed throughout the brain, which enables examination of the role of higher-order brain regions, such as those involved in attentional processes, while avoiding confounds in task-difficulty. In the present study, we utilised these advantages by assessing whether the RS FC of five visual RS networks (VRSN) – prior to training – is related to training success. We hypothesised that the strength of particular functional connections within these networks are associated with the attention-modulated increment in visual sensitivity (as previously described in Elshout et al., 2016), and thus to the patients’ attentional capacities.

To date, several mechanisms have already been proposed that can explain the variability in VRT success. For example, the “residual vision activation theory”, as proposed by Das and Huxlin (2010) or Sabel et al. (2011), describes that the extent of visual restitution depends on the presence and the amount of residual visual structures. By reactivating these structures restoration can be achieved. Some of these hypotheses have been confirmed by data, for example the prediction of VRT efficacy by relative visual defects (Poggel et al., 2004). Others, such as the presence of residual visual activity in primary visual cortex at locations without conscious vision (Papanikolaou et al., 2014), still await testing. For these reasons, we additionally identified areas of relative defects (areas of residual vision) in our patients and investigated how these relate to their training success. Specifically, we investigated whether the mechanism of visual restitution is related to the extent of residual vision. In line with previous literature (Guenther et al., 2009; Poggel et al., 2004; Sabel et al., 2011), we expect the size of the relative defect to be positively related to training outcome.

## 2. Materials and methods

The data described here are part of a larger project approved by the Central Committee on Research Involving Human Subjects Arnhem-Nijmegen in conjunction with the 1964 Declaration of Helsinki. Behavioural results have previously been reported in Elshout et al. (2016). In the present study, we focussed on static perimetry measurements (as measured with Humphreys Field Analyser (HFA)) and their changes (sensitivity in dB) during visual training in relation to prior-to-training RS functional MRI (fMRI).

### 2.1 Participants

We included thirty chronic (>10 months post-incident) post-chiasmatic stroke patients (15 males, age range: 26-69) with a homonymous visual field defect, whom gave written informed consent prior to participation. All patients had a macular sparing of >2 degrees, showed no signs of visual neglect, and were MRI eligible. Patients participated in two training rounds of each eight weeks, with in total at least 40 hours of training per hemifield. Static field perimetry (HFA 30-2 SITA Fast) measurements (i.e., sensitivity in dB) were obtained before training, after completion of the first and after completion of the second training round. Of the initial thirty patients, the HFA of five patients was considered unreliable due to a high rate of probe detection in the blind spot (>20%) and these patients were therefore excluded. Furthermore, two patients were excluded due to issues with the MRI data (see also Section 2.6), and three patients dropped out for personal reasons. Therefore, the final dataset consisted of data from 20 patients (11 right-sided, eight left-sided, and one bilateral VFD) (Table 1).

**TABLE 1.** Description of the patient sample

| No | Age<br>(y) | Sex | Type of field defect | Location of Lesion | Age of<br>lesion (mo) | Cause | Defect depth<br>(full field) |
| --- | --- | --- | --- | --- | --- | --- | --- |
| 1 | 61 | m | Scotoma | right occipital<br>cortex | 12 | Ischemic<br>stroke | 2.28 |
| 2 | 61 | m | Complete<br>Hemianopia | left occipital cortex | 34 | Ischemic<br>stroke | 14.75 |
| 3 | 56 | m | Quadrantanopia<br>(low) | left occipital and<br>parietal cortex | 18 | Ischemic<br>stroke | 7.61 |
| 4 | 69 | m | Incomplete<br>Hemianopia | left occipital cortex | 17 | Ischemic<br>stroke | 10.67 |
| 5 | 43 | f | Incomplete<br>Hemianopia | left occipital cortex | 20 | Ischemic<br>stroke | 10.46 |
| 6 | 57 | f | Complete<br>Hemianopia | left occipital cortex | 17 | Ischemic<br>stroke | 12.68 |
| 7 | 46 | f | Incomplete<br>Hemianopia | left occipital cortex | 22 | Ischemic<br>stroke | 13.92 |
| 8 | 52 | m | Incomplete<br>Hemianopia | right occipital<br>cortex | 29 | Ischemic<br>stroke | 12.83 |
| 9 | 33 | m | Complete<br>Hemianopia | left temporal cortex<br>and optic radiation | 18 | Hemorrhagic<br>stroke | 14.03 |
| 10 | 43 | f | Incomplete<br>Hemianopia | left occipital cortex | 21 | Ischemic<br>stroke | 10.71 |
| 11 | 43 | m | Bilateral incomplete<br>hemianopia | right and left<br>occipital cortex | 19 | Ischemic<br>stroke | 14.96 |
| 12 | 48 | m | Scotoma | right occipital<br>cortex | 17 | Ischemic<br>stroke | 6.01 |
| 13 | 53 | m | Incomplete<br>Hemianopia | right temporal and<br>parietal cortex<br>(optic radiation) | 17 | Ischemic<br>stroke | 7.97 |
| 14 | 39 | m | Scotoma | left occipital cortex | 85 | Hemorrhagic<br>stroke | 0.84 |
| 15 | 46 | f | Incomplete<br>Hemianopia | right occipital<br>cortex | 111 | Ischemic<br>stroke | 12.25 |
| 16 | 48 | m | Quadrantanopia<br>(up) | right occipital<br>cortex | 19 | Ischemic<br>stroke | 6.18 |
| 17 | 29 | m | Incomplete<br>Hemianopia | left occipital cortex | 21 | Ischemic<br>stroke | 9.05 |
| 18 | 49 | m | Complete<br>Hemianopia | left occipital and<br>temporal cortex | 38 | Hemorrhagic<br>stroke | 14.96 |
| 19 | 68 | m | Incomplete<br>Hemianopia | right occipital<br>cortex | 19 | Ischemic<br>stroke | 5.30 |
| 20 | 34 | m | Incomplete<br>Hemianopia | right occipital and<br>parietal cortex | 11 | Hemorrhagic<br>stroke | 8.62 |

### 2.2 Training paradigm

Following a randomised controlled cross-over and counterbalanced design, patients were trained successively at several locations within two predefined regions: one in the perimetric defect hemifield (primarily along the defect border) and one in the perimetric intact hemifield (at similar eccentricities as in the defect region). Thus, after two rounds, both regions had received training (see Figure 1 – A and B) and had alternately served as a control. During training, patients performed a visual discrimination task by directing spatially selective attention to the stimulus presented in the trained region. Specifically, patients had to covertly direct their attention, i.e. while suppressing eye-movements, towards the stimulus. The location of the training stimulus was primed by a straight line (in case of a Static Point stimulus) or indicated by the centre of a contracting flow field that filled the entire screen (in the case of the Optic Flow stimulus). In a previous paper on this patient cohort, Elshout and his colleagues (2016) reported no effect of training order (defect – intact, or vice versa) or training stimulus (static point or optic flow) on the training outcome. For a more detailed description of the training procedures, see Figure 1C and Elshout et al. (2016).

**FIGURE 1.**
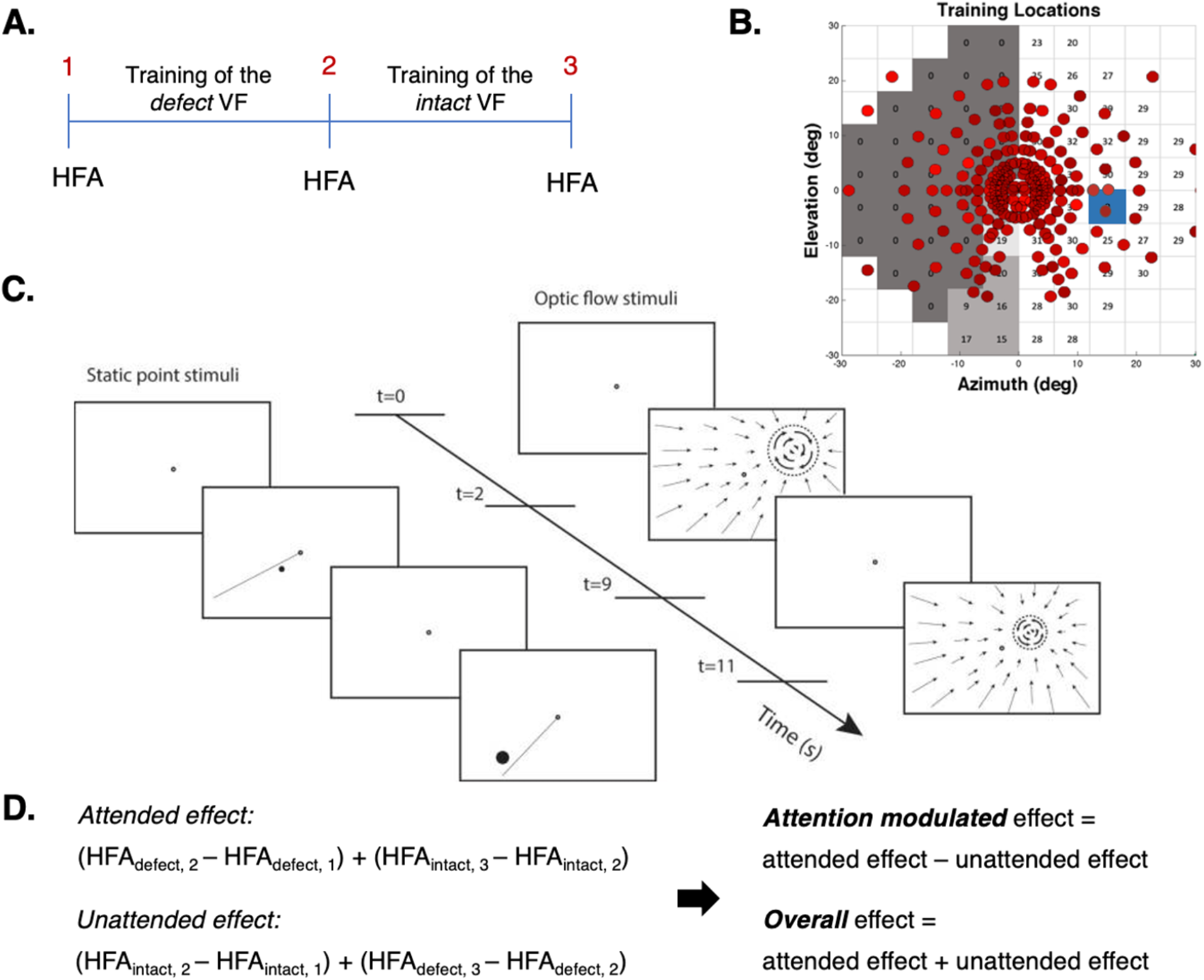
Training paradigm and procedure. **A.** Each patient participated in two successive training rounds: one with training stimuli in the defect VF and one with training stimuli in the intact VF. Panel A shows the training paradigm for half of the patients. The other patients started with training of the intact followed by a training of the defect VF. Because one hemifield was trained at a time, the other hemifield could serve as a control hemifield for that training round. The change in VF sensitivity was obtained monocularly, with static field perimetry (HFA, central 30 degrees), at three time-points: 1 (before training), 2 (after completion of round one) and 3 (after completion of round two). **B.** This panel shows the training locations, after both training rounds, presented on top of the visual field defect of one of the patients (patient no. 7, Elshout et al. (2016)). The training stimuli were presented at various locations within the visual field defect and, in the other training round, at similar eccentricities within the intact visual field. The dark grey patch marks the area of absolute, the lighter grey patch the area of relative and the lightest grey patch the area of minimal-no defect (see Section 2.3.2) at time point 1. The blue square represents the blind spot. For the training locations of all patients, see Elshout et al. (2016) (Supplementary Material). **C.** During each training round, patients performed a visual discrimination task with either a Static Point (left) or an Optic Flow (right) stimulus. First, patients were instructed to fixate on the centre of the screen (for 2s) as indicated by the black fixation dot. Then, the locus of the training stimulus was visually primed to the patient (for 7s) by either a straight line (in case of Static Point stimulus) or optic flow towards the patch (in case of the Optic Flow stimulus), followed by a fixation (for 2s). Lastly, the training stimulus was presented (for 7s) to which the patients had to direct spatially selective attention (covertly). Patients then had to decide on the location of the static point stimulus relative to the line or the rotation direction of the optic flow stimulus (Elshout et al. 2016). Used with permission). **D.** Change in sensitivity (dB) was evaluated for the trained and the control hemifield in both training rounds, and combined by category (trained or control), to assess attended and unattended training effects for the entire visual field. This enabled the computation of two VRT effects: an attention-modulated effect (i.e., the extent by which the attended training effect is larger than the unattended training effect) and an overall training effect (i.e., the sum of attended and unattended training effects). The panel shows the computation given the training order as presented in panel A.

### 2.3 Analysis of behavioural data

#### 2.3.1 Definition of the training effects

Following each training round a change in VF sensitivity, i.e., the mean decibel (dB) calculated from 38 measurement points (HFA, central 30 degrees), was determined for the trained and the control hemifield separately. This calculation was done relative to the previous measurement, not relative to the initial measurement, and thus quantified the effect of the latest training round only. From this, we calculated the overall sensitivity change in the attended (i.e., average change in the trained parts in successive training rounds) and the control (i.e., average change in the untrained parts). These two quantifications allowed us to examine two separate outcome measures: a training-specific effect (the attention-modulated training effect – i.e. the difference between the change in the attended and control hemifields) and the overall effect (the total change of both the attended and control hemifields). See Figure 1D.

Note that both outcome measurements characterise the visual sensitivity improvement by VRT for the entire visual field, rather than for the defective and intact visual field separately. In a previous report on this patient cohort, we showed similar training effects for the defect and intact visual field. Furthermore, for both the defect and intact side, larger visual improvements were found when trained (and thus attended) than when acting as the control area. For that reason, we considered their sum, which was significantly larger when attended than unattended (Elshout et al. 2016), as our training outcome parameter (see also Section 4.3 Full field improvements following VRT). Because we quantified our outcome measurements as sensitivity improvements over the entire visual field, we also took into account the defect depth (the average loss of sensitivity of the entire visual field) prior to training for each patient. This value indicates the patient’s maximum achievable improvement.

#### 2.3.2. Definition of defective subregions

To examine the relationship between the presence of residual vision and patients VRT outcome, we divided each patient’s defect visual hemifield (as measured at time point A) into three sub-areas: areas of absolute (<1 dB), relative (1-20 dB) or minimal-no defect (>20 dB), see also Figure 1B for an example. For each of these sub-areas, we calculated the size (i.e., the number of measurement points) and the proportion of points that showed an attention-modulated effect or an overall effect. The latter was used to evaluate where in the visual field the improvement was most apparent.

### 2.4 Image acquisition

MRI data were collected at the Donders Institute for Brain, Cognition and Behaviour using a SIEMENS MAGNETOM Skyra 3T scanner. Three high-resolution anatomical (T1) images were acquired just before, halfway through and immediately after completion of the training (MPRAGE sequence: voxel size = 1 mm isotropic, image matrix dimensions = 256*256*192, repetition time = 2300ms, echo time = 3.03ms, inversion time 1100ms, total acquisition time = 5min 21s). Resting-state blood oxygen level-dependent (BOLD) fMRI data were collected prior to training. A total of 1030 multi-echo images (voxel size = 3.5*3.5*3 mm, image matrix dimensions = 64*64*39, repetition time = 2000ms, flip angle = 80°) were obtained (interleaved sequence) at five different contrasts (echo times: 6.9, 16.17, 25.44, 34.71, and 43.98) (Poser et al, 2006). For two patients, the total number of volumes acquired was slightly lower (minimum of 917).

### 2.5 Pre-processing neuroimaging data

Neuroimaging data were analysed using Freesurfer (http://surfer.nmr.mgh.harvard.edu/), FSL (5.0.9; Jenkinson et al., 2012) and the ICA-AROMA package (Pruim et al., 2015). All preprocessing steps were performed at the individual level.

### 2.6 Anatomical pre-processing

A single anatomical scan was created by first aligning and then averaging three anatomical scans that were collected throughout the training (i.e., at time point 1, 2 and 3). The aligned and averaged scan was then structurally parcellated using Freesurfer to obtain a grey (GM), white matter (WM) and cerebrospinal fluid (CSF) mask. We used GM and WM masks to mask out the lesion site during registration and the subsequent analyses of the functional data.

### 2.7 Functional pre-processing

Multi-echo data were pre-processed using FSL by first applying motion correction and estimating the contrast-to-noise ratio (CNR) for each of contrasts before combining them in a CNR-weighted manner. Subsequent pre-processing steps included brain extraction, spatial smoothing (6mm) and non-linear registration of the anatomy scan to MNI (MNI_T1_2mm) with a warping resolution of 10mm. RS data were denoised using ICA-AROMA, a tool that identifies and removes head-motion-related artefacts. Further denoising was achieved by regressing out time series related to the mean activity in WM and CSF and by high-pass filtering (0.01 Hz). Lastly, denoised RS data were registered to MNI space (see also the top-left box of Figure 2). Multi-echo data of two patients could not be combined correctly due to unknown reasons, so these patients were excluded from further analyses.

**FIGURE 2.**
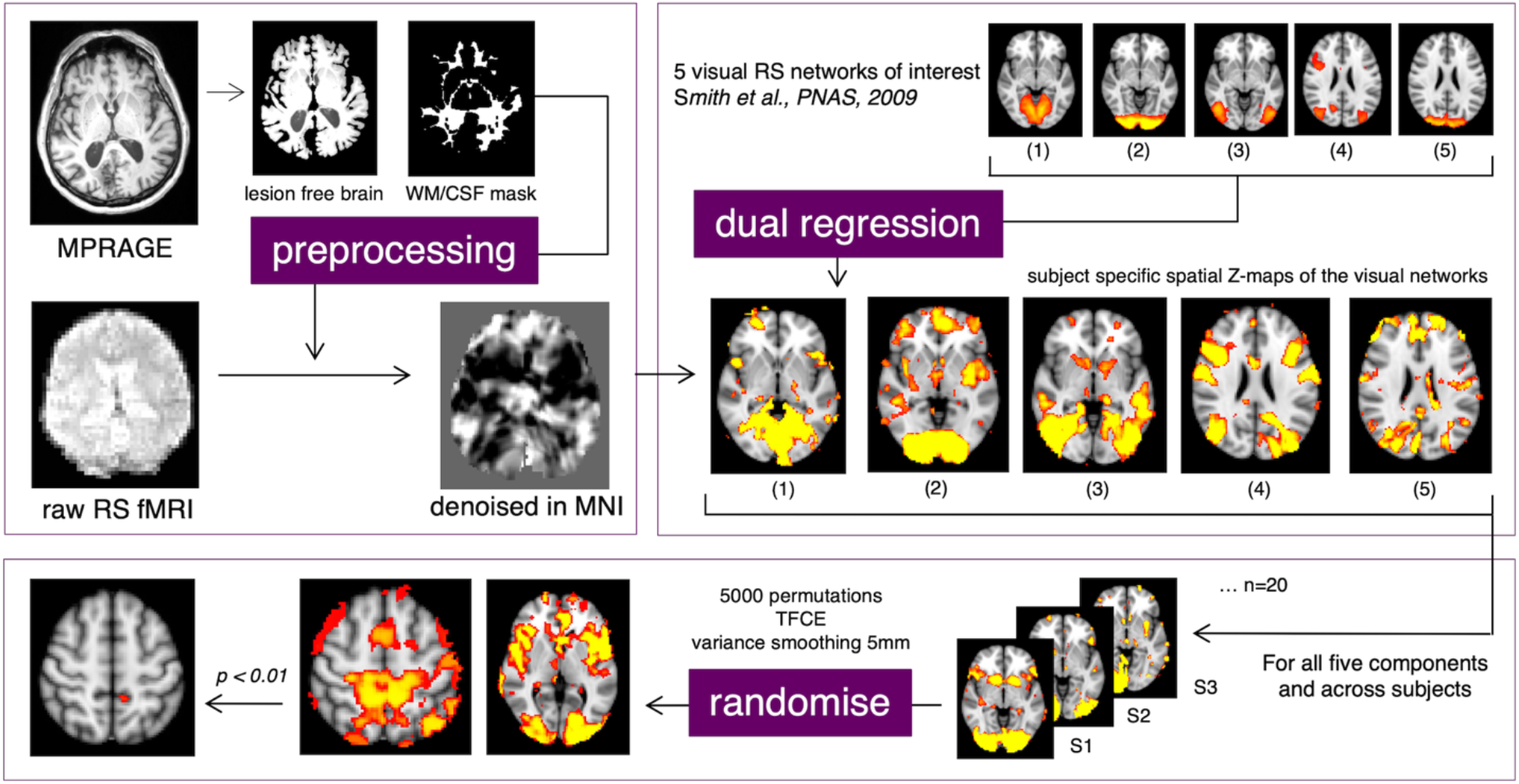
Processing pipeline. Top left: pre-processing steps of anatomical and functional data resulting in denoised functional data in MNI space. Top right: dual regression of 20 resting-state networks of interest resulting in patient-specific spatial maps of the networks. Only the VRSN of interest are depicted: 1) Medial Visual; 2) Occipital Pole; 3) Lateral Visual; 4) Oculomotor, and 5) Dorsal Visual Network. Bottom: permutation testing (using FSL randomise) of the VRSN, resulting in threshold-free cluster-enhanced t-statistics (Bonferroni corrected).

### 2.8 Generation of patient-specific spatial maps of resting-state networks

Because of the visual nature of the training, we focussed on five visual resting-state networks (VSRN) of interest that were selected from a set of 20 spatial maps of functional RS networks as described by Smith et al. (2009). These spatial maps are based on the RS-fMRI components of a 20-dimensional Independent Component Analysis on an independent dataset of healthy individuals. The VSRN of interest were: the Medial Visual, Occipital Pole, Lateral Visual (1_20_, 2_20_, 3_20_, respectively, as predefined by Smith et al. (2009)), and an Oculomotor (13_20_) and Dorsal Visual (16_20_) Network.

First, all 20 spatial maps were regressed onto the individual RS data, using dual regression (Beckmann et al., 2009), thereby generating patient-specific versions of these spatial maps and their associated time series. Specifically, for each patient, the 20 spatial maps were regressed (as spatial regressors in a multiple regression) onto the patient’s 4D space-time dataset. This resulted in a set of patient-specific time series for all 20 functional RS networks. Next, those time series were regressed (as temporal regressors, again in a multiple regression) onto the same 4D dataset. This resulted in a set of patient-specific spatial maps, one for each resting-state network, that contained Z-statistics. A voxel’s Z-statistic represents the standardised correlation of that particular voxel with the patient’s unique time series of a specific network. In other words, it describes how strongly the voxel covaries with the network and is thus related to the network. This will be referred to as the voxel’s *functional connectivity strength*. From this set of 20 spatial functional connectivity strength maps the five VRSN of interest were selected for further analyses (see top-right of Figure 2, Networks 1-5 respectively).

### 2.9 Randomised modelling

To test for a potential relationship between the patients’ spatial maps of FC and their training outcome, we performed nonparametric testing at group level. Specifically, we used a randomised modelling test (Winkler et al., 2014) based on 5000 permutations with threshold-free cluster-enhancement and variance smoothing (5 mm). In the model, we specified a training outcome (i.e., training-specific or overall effect) as the predictor, and the spatial maps of FC as the dependent variable. Furthermore, age and defect depth were included as covariates. For each map’s test outcome, a *p*-value < 0.01 (Bonferroni corrected) was considered significant, accounting for multiple testing of five networks of interest (see also bottom box Figure 2).

## 3. Results

### 3.1 Change in mean sensitivity of the entire visual field following VRT

The potential for visual sensitivity improvement varied considerably among patients: defect depth ranged from 0.84 dB to 14.96 dB with an average of 9.28 dB. The overall sensitivity change (mean = 0.92 dB, n=20) differed significantly from zero (one sample t-test: t = 5.00, df = 19, *p* < 0.001). Also, we found a significant training-specific effect (i.e., attended effect – unattended effect). Specifically, a significant difference of 0.43 db (one sample t-test: t = 2.12, df = 19, *p* = 0.047) was found between the contributions of the trained (mean = 0.67 dB, SE = 0.14) and the control hemifield (mean = 0.25 dB, SE = 0.13), see also Figure 3. No effect of lesion side (left or right) was found for either the training-specific (t = 1.11, df = 17, *p* = 0.28) or the overall effect (t = −0.02, df = 17, *p* = 0.99). One subject was excluded from this calculation due to the presence of bilateral cortical damage. Because we did not find an effect of lesion side on the training outcomes, it was not included as a confound in the subsequent analysis.

**FIGURE 3.**
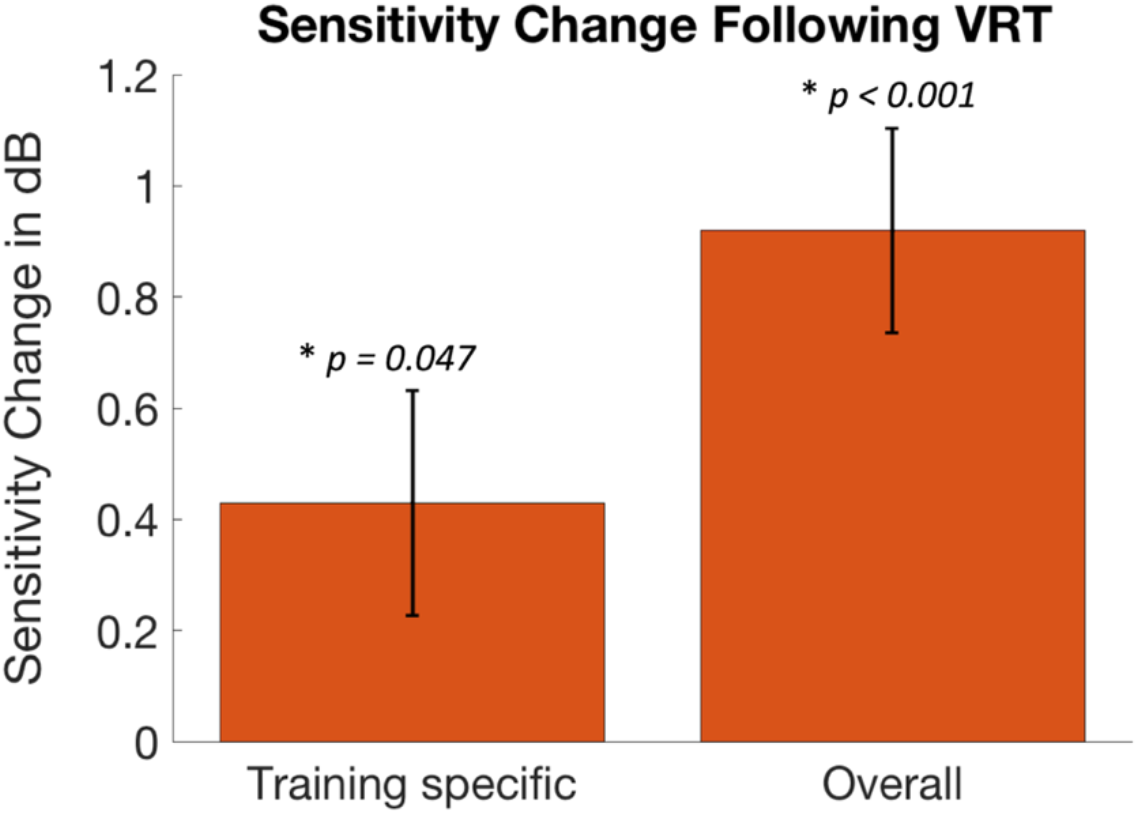
Sensitivity changes following VRT. Bars represent the training-specific (left) and the overall (right) change in mean sensitivity (in dB) of the visual field following VRT. Error bars represent Standard Error of the Mean. Significant improvements (*p < 0.05)* were found for both quantifications of the VRT effect.

### 3.2 Change in sensitivity following VRT per defective subregion

Table 2 shows the size (i.e., the number of HFA sample points) of the three sub-areas of the defective field (i.e., the absolute, relative and minimal-no defect), per participant at time point 1. Note that patient no. 11 has a bilateral visual field and therefore 76 measurements points and that patient no. 14 has only a minimal drop in sensitivity. Within brackets is the fraction of points that showed a training-specific (first value) and an overall (second value) effect, after completion of the VRT. The mean size of the absolute, relative and minimal-no defect were respectively 25.5, 4.9 and 9.7 measurement points. The relative and minimal-no defect regions showed larger proportions of measurement points with improved sensitivity (40-45%) compared to the absolute defect region (11-15%) (*Z* = 3.00, *p* = 0.003 and *Z* = 2.59, *p* = 0.010, respectively, Wilcoxon signed-rank test).

**TABLE 2.** Sizes of defective sub-areas. Bold value represents the size (i.e., the number of HFA sample points) per defect category. Values within brackets represent the fraction of points that showed a training specific (first value) and an overall (second value) training effect.

| No. | Absolute Defect<br>(proportion<br>improvement) | Relative Defect<br>(proportion<br>improvement) | Minimal – no<br>Defect (proportion<br>improvement) | No. of<br>Measurement<br>Points |
| --- | --- | --- | --- | --- |
| 1 | **11** (0, 0) | **7** (0.71, 0.57) | **20** (0.45, 0.35) | 38 |
| 2 | **37** (0, 0) | **1** (0, 0) | **0** (na) | 38 |
| 3 | **16** (0,0.06) | **7** (0.29, 0.57) | **15** (0.67, 0.80) | 38 |
| 4 | **24** (0.21, 0.21) | **4** (1, 0.75) | **10** (0.30, 0.80) | 38 |
| 5 | **26** (0.12, 0.15) | **2** (1, 0) | **10** (0.10, 0.50) | 38 |
| 6 | **32** (0.03, 0.03) | **5** (0, 0) | **1** (0, 0) | 38 |
| 7 | **28** (0.07, 0.11) | **8** (0.5, 0.5) | **2** (0, 1) | 38 |
| 8 | **31** (0.10, 0.13) | **7** (0.86, 0.71) | **0** (na) | 38 |
| 9 | **37** (0, 0.30) | **1** (0, 0) | **0** (na) | 38 |
| 10 | **34** (0.06, 0.06) | **2** (0, 0) | **2** 0, 0) | 38 |
| 11 | **56** (0.04, 0.09) | **8** (0.75, 0.75) | **12** (0.33, 0) | 76 |
| 12 | **6** (0.67, 0.50) | **16** (0.88, 0.75) | **16** (0.81, 0.50) | 38 |
| 13 | **11** (0.27, 0.55) | **16** (0.38, 0.56) | **11** (0.45, 0.36) | 38 |
| 14 | **0** (na) | **0** (na) | **38** (0.87, 0.87) | 38 |
| 15 | **32** (0.06, 0.06) | **0** (na) | **6** (0.17, 0) | 38 |
| 16 | **16** (0.06, 0.13) | **3** (0, 1) | **19** (0.63, 0.68) | 38 |
| 17 | **31** (0.06, 0.16) | **2** (1, 0.50) | **5** (0, 0.20) | 38 |
| 18 | **37** (0, 0) | **1** (0, 0) | **0** (na) | 38 |
| 19 | **27** (0.07, 0.04) | **2** (0.50, 0) | **9** (0.44, 0.22) | 38 |
| 20 | **18** (0.22, 0.22) | **6** (0.5, 0.50) | **14** (1, 0.71) | 38 |
| **Average** | **25.5** (0.11, 0.15) | **4.9** (0.46, 0.43) | **9.7** (0.39, 0.44) | na |

### 3.3 Neural correlates of the training effects

No significant correlation was found for the overall effect and any voxels’ FC strength with any of the VRSN tested (all *p* > 0.142). For the training-specific effect, however, we found a cluster of 18 voxels in the left hemisphere for which the FC strength with the Occipital Pole (OP) network was related to the training outcome (nonparametric permutation: t > 4.90, *p* < 0.01, family-wise error (FWE) corrected). This was a positive relationship: the stronger the FC strength with the OP network, the larger the training-specific change in mean sensitivity (henceforth referred to as the attention-modulated training effect, see Figure 4B). Although associated with the OP network, comprising primarily the Occipital Pole area, the cluster was located at the anterior division of the left Precuneus (Superior Parietal Lobule area 5M, Juelich Histological Atlas; Eickhoff et al., 2005; Scheperjans et al., 2008). This indicates that brain regions outside the main core of the network could still contribute to a network’s function connectivity. The peak of the effect, the voxel with the max t-statistic (t = 5.56, *p* = 0.0082), was found at [−10 −44 54] MNI see also Figure 4A – left hemisphere. No such relationship to the attention-modulated training effect was found for any of the other VRSN (all *p* > 0.089).

**FIGURE 4.**
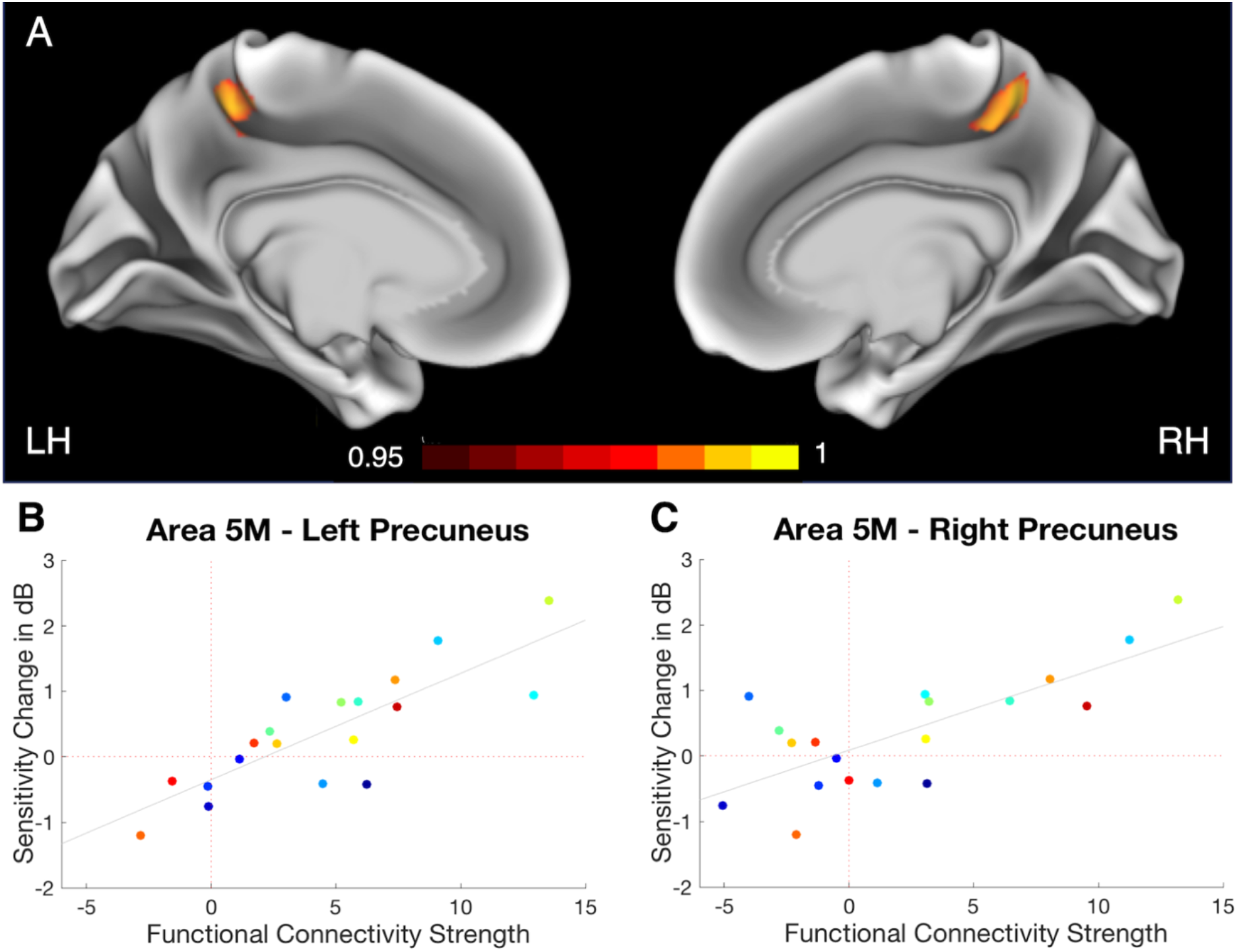
A | Results of resting-state fMRI analysis projected on the inflated brain. Maps depicted in red/yellow represent the randomised results (1-p maps, *p* < 0.05) for the relationship between the attention-modulated training effect and the voxel’s FC strength with the OP network for the left (LH) and right (RH) hemisphere. We found a significant cluster (*p* < 0.01) in the anterior division of the left Precuneus, area 5M (core: [−10 −44 54] MNI), and a marginally significant cluster (*p* < 0.05) in the anterior division of the right Precuneus, again area 5M (core: [11 −46 54] MNI). **– B and C | Relationship between attention-modulated training effect and FC strength.** Data is presented for left and right 5M separately after the removal of one patient who’s right 5M fell within his/her lesion site. Each data point represents one patient, points of the same colour represent the same patient. Mean sensitivity change is presented for the attention-modulated training effect and in relation to the patients’ max FC strength of area 5M with the OP network. As the FC becomes stronger, the attention-modulated training effect becomes larger.

### 3.4 Bilateral neural correlates

Although the significant cluster found in the Precuneus was unilateral, we found an additional marginally significant cluster (*p* < 0.05) of 285 voxels in the anterior division of the right Precuneus (see Figure 4A – right hemisphere). Inspection of the individual lesion sites revealed that this cluster partially (for ~ 21%) fell in the lesion area of one patient (see Supplementary Material). A posthoc analysis (randomised modelling including the attention-modulated training effect and the OP network) with this patient excluded revealed an enlarged cluster in the left Precuneus (core: [−8 −44 54] MNI, size = 58 voxels; randomise permutation: t > 4.75, *p* < 0.01, FWE corrected) and an additional cluster in the right Precuneus (core: [10 −46 56] MNI, size = 26 voxels; randomise permutation: t > 4.88, *p* < 0.01, FWE corrected). See also Figure 4 B and C for the visualisation of the relationship between the OP network functional connectivity strength for the left and right 5M separately and the patients’ attention-modulated training outcome.

### 3.5 Non-visual RS networks

In this study, we examined only the FC profiles within VRSN (as described by Smith et al., 2009), as we expected the training effects to result primarily from a change in processes related to visual perception. To determine whether improvements could also be related to processes with primarily attentional or non-visual sensory components, we repeated the analyses for the non-visual networks of interest. Specifically, we tested three attentional networks (the left and right Fronto-Parietal (9_20_ and 10_20_, resp.) and the Dorsal Parietal (12_20_), the Sensorimotor (6_20_) and the Auditory (7_20_) network, as described by Smith et al., 2009. No significant relationship with either of the VRT effects was found for any of these networks (all *p* > 0.06).

### 3.6 Correlation of the defect size with training effects and the functional connectivity of the Precuneus

For both training effects, larger proportions of measurement points with improved sensitivity were found for the relative and minimal-no defect areas compared to the absolute defect area (see also section 3.2). To further explore this relationship, we correlated the sizes of the three sub-areas with both training effects and the functional connectivity strength of the Precuneus with the OP network. Neither the overall training effect correlated significantly with the size of any of the sub-areas (absolute defect: *r* = −0.36, *p* = 0.14; relative defect: *r* = 0.18, *p* = 0.44; minimal-no defect: *r* = 0.41, *p* = 0.09), nor the attention-modulated training effect (absolute defect: *r* = 0.03, *p* = 0.91; relative defect: *r* = 0.38, *p* = 0.12; minimal-no defect: *r* = −0.06, *p* = 0.81). The functional connectivity strength of the Precuneus correlated only marginally significant with the size of the relative defect (*r* = 0.48, *p* = 0.059), and not with the other two sub-areas (absolute defect: *r* = −0.14, *p* = 0.55; minimal-no defect: *r* = 0.01, *p* = 0.98).

## 4. Discussion

Our main finding is that RS FC between the Precuneus and the OP network is related to the attention-modulated component of VRT-induced VF improvements in hemianopia. Specifically, the stronger the functional connection is prior to training, the more likely it was that the patient benefitted from the training. This finding builds on the result that patients benefitted from VRT discrimination training at known (primed) locations by improved processing of spatially unpredictable visual events in parts of the VF (Elshout et al., 2016). In other words, training success does not rely on knowledge of the stimulated location, as offered during the training, but rather on the patients’ more general improvement in their ability to direct attention to visual stimuli (as indicated by detection of stimuli at *unknown* locations during perimetry). These results suggest that the Precuneus plays a defining role in this success by mediating the attentional processes. Consequently, our results suggest that the RS FC strength of the Precuneus with the OP network could be used as a biomarker to identify hemianopia patients with a potentially high training yield. This is useful because VRT is laborious, time-consuming and costly, and not all patients benefit equally. Our study thus provides both new insights and a novel avenue towards probing the neural origin of the variation in VRT outcome, which can ultimately contribute to improving the success rate of VRT.

### 4.1 A functional neural correlate of the attention-modulated training effect

We found a cluster of voxels in the anterior division of the left Precuneus, and a marginally significant cluster in the anterior division of the right Precuneus (both area 5M), whose FC strength with the OP network was positively related to the magnitude of the attention-modulated training effect. In other words, the more strongly these areas were connected to the OP network, the larger the attention-modulated visual improvement became. No such relationship was found for any of the other visual networks tested or for the overall training effect. The latter observation suggests that this particular FC is not predictive of a general improvement of the VF. Instead, the finding relates to the specific effect of selective spatial attention, which is necessary for performing the training well, on the visual improvements. It suggests that the engagement of areas 5M in the OP network could be a modulator of VRT outcome by targeting such attentional processes.

Our findings complement our current understanding of the variability of VRT efficacy by presenting a new essential functional and attention related component. Poggel et al. (2004) previously showed that attentional processes play an essential role in visual restitution. Specifically, they showed that attentional cueing of areas of relative defects (or residual vision) boosted the treatment outcome of VRT in patients with visual field defects. Our study showed an improvement of visual detection by specifically training directing spatial attention, which extent was strongly related to the functional network connectivity of the Precuneus.

Other studies have suggested that the mechanism of visual restitution involves activation of residual vision. For example, Sabel et al. (2011) described how visual structures with residual visual functioning could, to a certain extent, be reactivated or restored depending on its size and its residual activation. These structures include, for example, areas of partial damage in the visual cortex (or areas of relative defects). Previous studies have indeed shown visual improvement after repetitive stimulation of these structures (Poggel et al., 2004; Sabel et al., 2011). Likewise, Papanikolaou et al. (2014) observed dissociations between the perceptual and retinotopic maps in patients with homonymous VFDs. Specifically, in a subset of patients, they found V1 visual field coverage, as measured by fMRI, to be larger than predicted based on perimetry maps. They attributed this difference to the presence of residual vision of “spared islands” in V1 that are responsive, but whose responses are not sufficient to achieve visual awareness. Furthermore, they also suggested that the presence and extent of these spared regions may reflect the capacity of V1 to recover function and could, therefore, contribute to training-induced improvements. Indeed, it has been shown that hemianopia patients can improve in parts of their absolute defect (Das and Huxlin, 2010b; Huxlin et al., 2009), but none of these studies collected data enabling testing for a possible relation between training-induced improvements and neural responses to stimulation inside the perimetric scotoma. In our study, we did not find a statistically significant relationship between the size of the relative defect (i.e., the area of residual vision) and either the attention-modulated or the overall training effect. This means that for our cohort, larger relative defects did not result in better VRT outcome. We did, however, observe larger proportions of measurement points with improved sensitivity within the areas of relative defect compared to the areas of absolute defect. Furthermore, we did find a trend for a positive relationship between the size of the areas of relative defect (and not the areas of absolute or the minimal-no defect) with the functional connectivity strength of the Precuneus with the OP network. One could argue that, because of this relationship, the connectivity strength of the Precuneus is primarily related to the activation of the areas of residual vision and not to attention. However, we argue that if indeed areas with a relative defect evoke a stronger OP network connectivity of the Precuneus, and the latter is thus independent of attentional modulation, we would expect to find such Precuneus network state for the overall training effect as well but which was not the case. Instead, we suggest that, since visual restitution was more apparent within the relative area of defect, the Precuneus may mediate attentional modulation more effectively in this area. It is important to point out that here we considered only the size of the relative defect, as examined by the number of measurement points with a sensitivity of 1-20db. It is possible that other characterizations of the size of the relative defect, as described by Guenther et al. (2009), play a role in training efficacy, in addition to the Precuneus network connectivity.

### 4.2 The anterior division of the Precuneus: a modulator of attentional capacities?

The Precuneus, the posterior region of the superior parietal lobe, is known to play an essential role in the implementation of a wide range of higher-order cognitive functions. It is believed to aid processes like visuospatial attention and updating (Le et al., 1998; Mahayana et al., 2014; Müller et al., 2018; Nagahama et al., 1999), and to modulate conscious processes (Cavanna, 2007; Cavanna and Trimble, 2006; Kjaer et al., 2001; Vogt and Laureys, 2005). Furthermore, it has been reported to play a role in tasks involved in the control of covert attention (Han et al., 2003; Mao et al., 2007; Sali et al., 2016). The Precuneus has also been associated with saccadic eye movements (Berman et al., 1999; Luna, 1998; Petit & Haxby, 1999), a process tightly linked to attention allocation. In particular, it has been reported to be involved in anti-saccades, i.e. voluntary eye movements during which reflexive saccades towards a peripheral visual cue are suppressed and directed towards the mirrored cue location in the opposite hemifield (Brown et al., 2006; Brown et al., 2007; Dyckman et al., 2007; Kimmig et al., 2001). While saccades are not essential for the control of attention, attention does play a crucial role in the control of saccades (Zhao et al., 2012). Furthermore, preparatory pre-saccadic attention shifts facilitate visual processing at the saccadic locations (Baldauf and Deubel, 2008; Deubel and Schneider, 1996; Jonikaitis and Deubel, 2011). Through covert attention, the visual system can make predictions about information present in the periphery, which in turn boosts the quality of visual representations. This process is also referred to as predictive remapping, the dynamics in receptive fields during saccade preparation (Carrasco and McElree, 2001; Findlay, 2005; Harrison et al., 2012; Mathôt and Theeuwes, 2010; Ritchie et al., 2012; Rolfs et al., 2011; Wolfe and Whitney, 2014; Zhao et al., 2012).

In our study, the ability to suppress eye movements was essential to perform the visual discrimination task correctly. At the same time, a successful covert attention shift towards the stimulus in the periphery helped the patient to perform the task well. We hypothesise that the above-mentioned saccadic processes may have boosted these attentional operations. In line with its known role in preparatory saccadic processes, the Precuneus may mediate the awareness recovery processes. In other words, a strong functional link between the Precuneus and the OP network may enhance preparatory saccadic behaviour which, in turn, facilitates visual awareness. Although the OP network concerns the foveal representation of the visual field, the visual field improvements were primarily found in its periphery. The functional association of the Precuneus with the OP network, but not with any of the other networks, may be related to the underlying predictive remapping process, as discussed above, where peripheral information is fed back to the cortical areas representing the fovea around the time of a saccade. Interestingly such remapping is known to raise awareness of flashed targets in the patient’s VF defect that disappear before the saccade brings that target in the intact VF (Ritchie et al 2012).

### 4.3 Full field improvement following VRT

In this study, we reported on visual improvements of the full VF rather than solely its defect parts for which it seems most likely that clinically relevant improvements will be obtained. Interestingly, a previous report on the same cohort of patients showed similar improvements for both the perimetric defect and intact VF (Elshout et al., 2016), which suggest that also in the intact parts of the VF clinically relevant improvements can be obtained. This complements various reports that show perceptual deficits in the “intact” parts of the VF (for review: Bola et al., 2013; Cavézian et al., 2015; Clatworthy et al., 2013; Geuzebroek and van den Berg, 2017). Furthermore, is has been suggested that this observation partially accounts for the subjective visual impairment as experienced by the patients and should therefore also be considered when examining VRT effects (Bola et al., 2013).

### 4.4 The controversy of VRT efficacy

VRT has been criticised due to inconsistent outcomes in previous research, ranging from significant improvements to minimal or no effects at all. A source of this inter-subject variation could be that the perimetric techniques are subjective and biased towards a positive outcome and that VRT outcomes are confounded by eye movements (Pouget et al., 2012; Reinhard et al., 2005). Indeed, the reliability of an extension of the visual field from perimetric data can be hampered by insufficient eye fixation control. In our study, however, we used Humphrey’s perimetry to assess sensitivity improvement of the visual field and rejected any outcome even if only one of the perimetric measurements showed insufficient fixation stability. Furthermore, Kasten et al. (2006) studied the potential confounding aspect of eye movements in VRT success and showed it to be independent of eye-movements. This observation has been confirmed by later studies on visual field training that controlled for eye-movements (Bergsma and Van Der Wildt, 2010).

Another relevant source of variation is the size of the individual’s defect or relative defect (Gall et al., 2016; Poggel et al., 2004; Sabel et al., 2011). In a systematic review on VRT efficacy, Bouwmeester et al. (2007) reported that the majority of the studies did not take the size of the VFD into account when evaluating the visual improvement following VRT. In our study, defect depth was included in the analysis as a covariate, accounting for individual differences in training outcome caused by variations in the degree of their defect. Furthermore, we found no significant correlation between the extent of the relative defect and any of the training outcomes.

We argue that our results cannot be explained by the confounding effect of eye movements or by the differences in sizes of the (relative) visual field defects. Instead, our results showed that the inter-subject variation can be explained the patients’ attentional capacities and which are modulated by the Precuneus’ functional network connectivity.

### 4.5 Clinical implications

For the patients, participation in extensive VRT programs (ranging from weeks to months) can be experienced as quite exhausting. Furthermore, objective visual improvements, even when substantial, might not be perceived subjectively as improvements. In another study, we evaluated the improvement in personal activities of daily living as measured with the Goal Attainment Scale (GAS) for a larger cohort of patients (including those presented here). We found a relationship between objective visual improvements after training, as assessed by Goldmann perimetry and the GAS scores. Furthermore, for the Humphrey perimetry, we found a linear relationship only between GAS scores and the directed training effects for the full visual field (Elshout et al., 2018). These findings emphasise the value of our quantification of the change in visual field sensitivity, i.e. as a full field measure, since it is relevant to the visual field used in daily life activities (as reflected with GAS). Other studies have also reported subjective improvements, after evaluating patients’ scores on Quality of Life and Activities of Daily Life assessments (Gall et al., 2010; Mueller et al., 2007; Poggel et al., 2004; Poggel et al., 2010; Sabel et al., 2004). These findings highlight the clinical relevance of VRT training.

No or a minimal gain after an extensive period of training can be disappointing for the patients. To minimise such undesired outcomes, appropriate identification of patients with high training potential prior to training participation is warranted. We acknowledge that the use of RS fMRI as a screening tool is not the most obvious, due to its high financial costs and the fact that not every patient will be MRI eligible. Furthermore, we used an extensive scanning protocol (i.e., about 30 minutes of scanning), which may not always be practicable to follow. From such a practical point of view, it would be easier to predict treatment outcome based on features from perimetric charts that describe the loss of visual function (see also Guenther et al., 2009, Gall et al., 2013) or behavioural outcome measures. Yet, these will not be able to quantify a patient’s potential training success. Neither does it add to our understanding of the underlying neural mechanism of the effect of attentional processes on VRT outcome. Therefore, we emphatically advocate the further exploration (see also Section 4.6 Future directions) of our finding of a functional neural correlate of VRT outcome and its potential as a biomarker to indicate a patient’s potential to direct spatial attention. It contributes to our understanding of the inter-subject variability in VRT efficacy, and may, therefore, serve as an indicator for VRT success. Furthermore, to date, no objective or alternative screening tools are available.

### 4.6 Future directions

Due to the nature of the study, we identified a neural correlate for an attention-modulated effect of VRT but were unable to make firm statements about the predictive capacity of the Precuneus’ RS FC of VRT effect. To validate the FC of the Precuneus as a predictor of training outcome, more testing is needed. For that reason, a prediction study, using a new cohort of hemianopia patients who are scheduled for training, is being planned by several of the current study’s authors.

## 5. Conclusion

Our study presents a potential functional mechanism that underlies the effect of attention on visual restitution. We argue that the engagement of the Precuneus may modulate patients’ attentional capacities. Specifically, we found that a strong RS FC of the Precuneus with the OP network prior to VRT reflects the training-induced visual improvement associated with the direction of spatially selective attention. Due to its role in saccadic performance, we speculate that the Precuneus mediates preparatory saccadic behaviour with enhanced visual processing as a result. The strength of this RS FC of the Precuneus in chronic hemianopia patients may serve as an imaging-based biomarker, predictive of capacity to direct spatial attention. Thereby, this biomarker may help identify patients with a potentially high training yield.

## Abbreviations

FC: functional connectivity
fMRI: functional MRI
FWE: family-wise error
GAS: goal attainment score
GM: grey matter
HFA: Humphreys field analyser
RS: resting-state
VF: visual field
VFD: visual field defect
VRSN: visual resting-state networks
VRT: visual restitution training
WM: white matter

## Supplementary Material

Each patient’s lesion site was localised using the lesion_gnb toolbox for SPM (Griffis et al., 2016). Of the 20 patients, eight had a lesion in the right hemisphere, and 11 had a lesion in the left hemisphere. One patient had a bilateral lesion. Investigating the overlap between the significant cluster in the left Precuneus and the patients’ lesion site showed that the lesion site did not cover the left Precuneus. One patient showed overlap between his or her lesion site and the marginally significant cluster in the right hemisphere. More specifically, in this cluster (285 voxels in total), 60 voxels fell within the patient’s lesion site. Figure I shows the marginally significant cluster in the right hemisphere projected on the anatomical image of this patient’s brain.

**FIGURE I.**
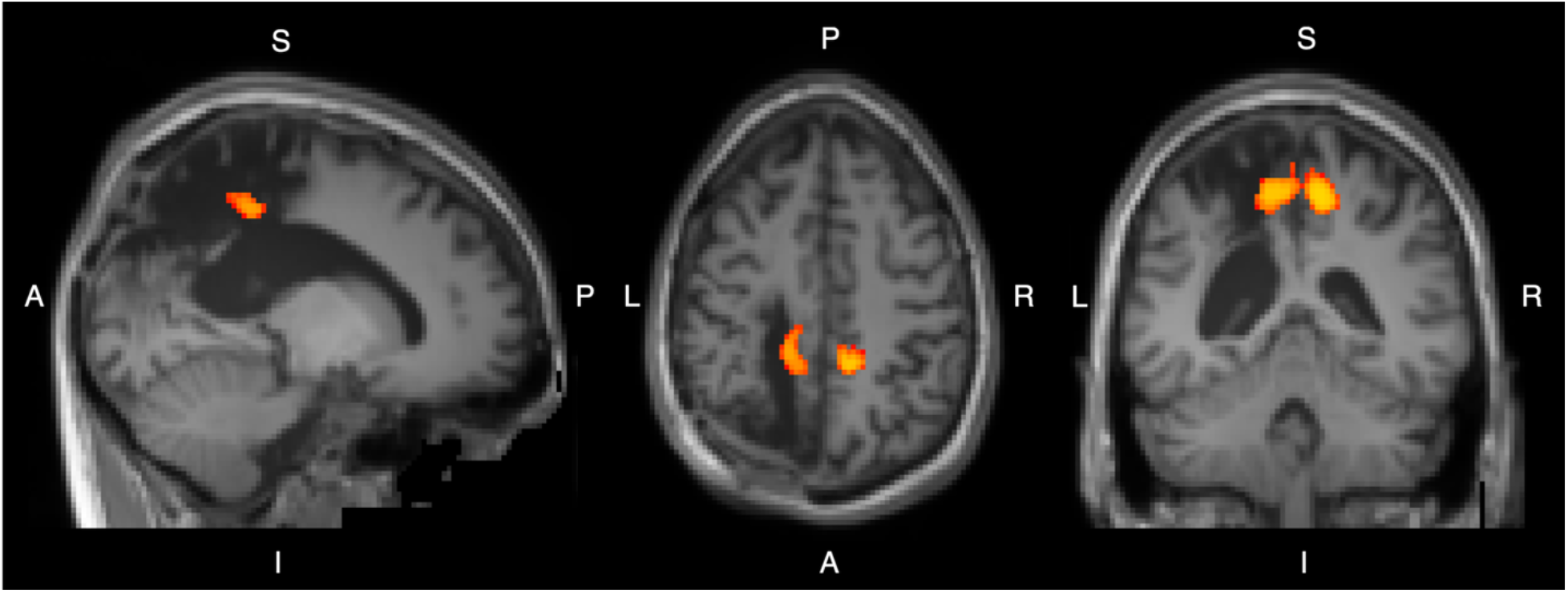
Randomised results partially overlap lesion site in one patient. The randomise results of the Occipital Pole network and the obtained attention-modulated change in sensitivity, thresholded at *p* < 0.05, projected on the anatomical image of one of the patients (normalised to MNI). Note that the marginally significant cluster in the right hemisphere falls within the patient’s lesion site. From the anatomical image identifying information is removed using mri_deface, the automated defacing tool of Freesurfer (http://surfer.nmr.mgh.harvard.edu/fswiki/mri_deface). Clinical data is used for publication with the permission of the patient.

## Acknowledgements

HNH was supported by the Research Institute of Brain and Cognition of the Graduate School of Medical Sciences, University Medical Center Groningen, the Netherlands. HNH and FWC have been supported by the European Union’s Horizon 2020 research and innovation programme under the Marie Sklodowska-Curie grant agreement No 641805 and No 675033). KVH gratefully acknowledges funding from the Netherlands Organisation for Scientific Research (NWO VENI Grant Project no. 016.Veni.171.068). Resting-state data were collected as part of the NWO ZonMW-Inzicht Grant Project no. 94309003 to AvB. The authors wish to thank all the patients for taking part in this study. Furthermore, we appreciate the general support of Paul Gaalman of the Donders Institute during the preparation and scanning of the patients, and Freekje van Asten for performing the Humphreys Perimetry on the patients.

## Data availability statement

The data that support the findings of this study are available from the corresponding author upon request.

**Declarations of interest: none**

